# Topslam: Waddington Landscape Recovery for Single Cell Experiments

**DOI:** 10.1101/057778

**Authors:** Max Zwiessele, Neil D Lawrence

**Affiliations:** University of Sheffield, Department of Computerscience

## Abstract

We present an approach to estimating the nature of the Waddington (or epigenetic) landscape that underlies a population of individual cells. Through exploiting high resolution single cell transcription experiments we show that cells can be located on a landscape that reflects their differentiated nature.

Our approach makes use of probabilistic non-linear dimensionality reduction that respects the topology of our estimated epigenetic landscape. In simulation studies and analyses of real data we show that the approach, known as topslam, outperforms previous attempts to understand the differentiation landscape.

Hereby, the novelty of our approach lies in the correction of distances *before* extracting ordering information. This gives the advantage over other attempts, which have to correct for extracted time lines by post processing or additional data.

## 1 Introduction

High-throughput single-cell real-time polymerase chain reaction gene expression measurements (Section S2) are new and promising techniques to give insights into the heterogeneous development of individual cells in organism tissues [19]. However, interpretation of measurements can be highly challenging.

Waddington [33,34] proposed a representation for understanding the process of differentiation, known as Waddington’s landscape or the *epigenetic landscape.* The idea is that differentiated cells are located at different points on the epigenetic landscape with particular paths through the landscape more likely than others due to its underlying topology (Figure 1). Originally, this landscape represents the quasi-potential function of genetic network dynamics and is shaped by evolution through mutational re-wiring of regulatory interactions [16]. In this context, the landscape is created by a complex set of interactions between transcription factors, genes and epigenomic modifications. Unpicking the mechanism behind this relationship is extremely challenging [2,16,20,35,36]. Instead we propose an alternative, data driven approach based on machine learning algorithms and judicious application of probabilistic methods. In this paper we reconstruct such landscapes from rich phenotype information through probabilistic dimensionality reduction. In particular, we extract maps of the epigenetic landscape given the observations of *gene expression.* The mathematical underpinnings of mapping involve a projection from a low dimensional space to a higher dimensional space. Classically we might wish to project the three dimensional world around us down to two dimensions for use as a map or a chart. Formally this involves a mapping, **f**(·) from the positions in the two dimensional space, **x**, to our measurements, **y**:

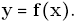

**Figure 1.**
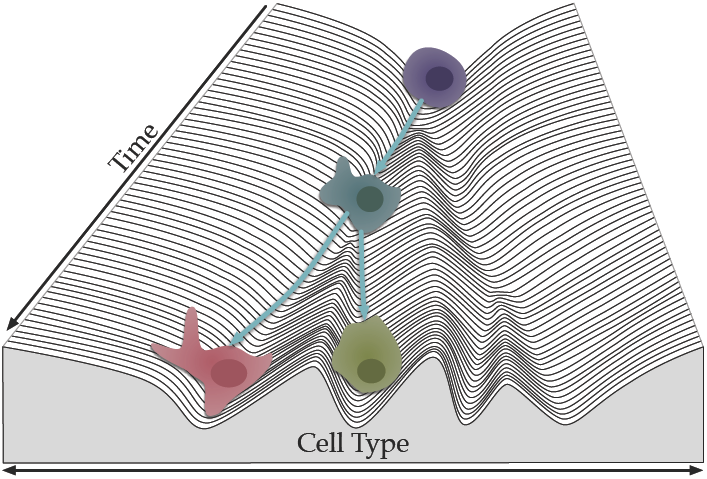
Waddington landscape representation for differentiating cells. The topology of the landscape is created by genetic network dynamics, shaped by evolution through mutational re-wiring of regulatory interactions. The cells follow along the topology – stochastically deciding at key junctions for differentiation paths.

In epigenetic landscapes, rather than considering the high dimensional measurements to be direct measurements of the physical world around us, we instead observe a rich phenotype, such as the gene expression of an individual cell, **y**. Our aim is to develop a coherent map such that the position of each cell, **x**, is consistent with cells that are expressing a similar phenotype. In other words, if two cells have a similar gene expression they should be located near to each other in the map, just as two people located near to each other in a real landscape would have a similar view.

The utility of a map is given by the precision in which it can be recreated. Geographic surveys were originally created through triangulation and laborious ground level surveys. The challenges we face for the epigenetic landscape are somewhat greater. In particular the measurements of phenotype are subject to a great deal of noise, particularly in single cell experiments, in other words there is a mistiness to the observations. Further, we cannot access all areas. We can only query individual cells as to their particular phenotype, we cannot move around the landscape at will. Finally, there is a complex, most likely non-linear relationship between any location on the map. Thus, we have to estimate the full smooth map and its distortions from the discrete observed points in form of cells and their gene expression patterns.

We are inspired by challenges in robotics: in robot navigation a robot facing a landscape for the first time needs to continually assess its current position (the values of **x**) and simultaneously update its estimate of the map (the function *f*(·)). This challenge is known as as simultaneous localisation and mapping (SLAM [27]).

For example Ferris *et al.* [9] showed how simultaneous localisation and mapping could be formed by measuring the relative strength of different WiFi access points as it moves around a building. When you are near to a given access point you will receive a strong signal, when far, a weak signal. If two robots both perceive a particular access point to have a strong signal they are likely to be near each other. We can think of the WiFi access points as landmarks. In our case landmarks are the (noisy) gene expression measurements. If two cells have a similar set of gene expression measurements they are also likely to be near each other. A further challenge for our algorithm is that gene expression measurements are very high dimensional and can be extremely noisy. Because of the analogy to SLAM algorithms and our use of topology to develop the landscape we refer to our approach as *topslam* (topologically aware simultaneous localisation and mapping).

Quantitative determination of single-cell gene expression is commonly used to determine the—known to be heterogeneous—differentiation process of cells in cancer [7] or in the early development of multicell organisms [14]. The measurement of single cells, however, can give rise to systematically introduced errors in the identification of sub processes in the cell and in the assignment of cells to their specific cell-lines. This is due to the low amounts of mRNA available in single cells: the mRNA requires amplification using polymerase chain reaction (PCR, see e.g. [11,15,21]).

These technical limitations complicate analysis: they introduce non-linear effects and systematic errors. So as promising as high throughput methods are, they require sophisticated analyses to resolve confounding factors. By providing the scientist with the underlying Waddington landscape for cells in a given experiment, along with the location of each cell in the landscape, we hope to significantly simplify this process. Unpicking the nature of the genetic landmarks in the presence of noise typically exploits *feature extraction,* where high dimensional gene expression data has its dimensionality reduced [10,14,23,24], often through linear techniques. However, it is difficult to determine the number of dimensions to use for further analyses [3-5].

### 1.1 Dimensionality Reduction

Dimensionality reduction gives a view on the landscape of the underlying biological system. To perform dimensionality reduction we need a mathematical model that extracts the salient aspects of the data without exhibiting vulnerability to confounding factors such as technical or biological noise.

Probabilistic models aim to trade off the useful structure with the confounding variation through specifying probability distributions for each component. We consider non-linear probabilistic models that not only model the landscape as a non-linear surface (think of an irregular skiing piste, in which you want to turn into the flat bits, as opposed to a flat beginners slope, where you can just go in a straight line), but also allow us to determine the dimensionality necessary to explain the gene expression variation, while explaining away the noise through a separate model component.

Linear methods can also be given probabilistic underpinnings, but they suffer from the severe constraint of only allowing the landscape to be linearly related to the genetic landmarks. Conversely deterministic (i.e. non-probabilistic) non-linear methods do not offer a principled approach to separating the confounding variation from the landscape’s underlying structure. It can be hard to grasp topographical relationships due to the deterministic nature of the technique. Either additional data or additional correctional deterministic algorithms are necessary for a coherent mapping [1,25].

We make use of the *Bayesian Gaussian process latent variable model* (Bayesian GPLVM [29]), a probabilistic dimensionality reduction technique that extracts the relevant dimensionality of the latent embedding as well as expressing a non-linear model. Further, we make use of the *geometry* of the underlying map by exploiting recent advances in metrics for *probabilistic* geometries [30].

### 1.2 PCA and Graph Maps

An approach such as principal component analysis (PCA) makes an assumption of a *linear* relationship between the high dimensional measurements and the cell’s location in the landscape. This limiting assumption is normally alleviated by proceeding in a two step manner. First PCA is done for all data, then the locations in the linear map are clustered and a further PCA is applied to each cluster separately, giving one coordinate system per cluster [14] (see also [8,28] for an elegant implementation of this approach and more).

Islam *et al.* [18] developed a graph based method, using similarities of cell profiles to characterise two different cell types in a so called “graph map”. Linear cell-to-cell correlation is used to create a 5 nearest neighbour graph. The graph is then visually adjusted by a force-directed layout to visualise cell-to-cell correlations. In topslam we only make use of a graph to extract shortest distance between cells and not for visualisation.

The Waddington’s landscape [33,34] can be seen as a non-linear map for the branching process of cells, where the cell process is described as a ball rolling down a hill following stochastically (by e.g. cell stage distribution) the valleys of the hillside (Fig. 3).

For topslam, the underlying probabilistic dimensionality reduction technique (Gaussian process latent variable model) has been successfully used in other applications to single cell transcriptomics data, e.g. for visualisation [3], to uncover sub populations of cells [4] and to uncover transcriptional networks in blood stem cells [22].

The novelty of our approach is to not correct *after* extraction of graph information, but to correct the distances the graph extraction *uses to extract* information. We can do that by estimating the underlying Waddington landscape along differentiation of cells.

### 1.3 Independent Component Analysis and Non-linear Dimensionality Reduction

Recovery of the epigenetic landscape as an intermediate step facilitates the extraction of other characteristics of interest, such as pseudo time, in cell stage development. For example Trapnell *et al.* [31] apply independent component analysis (ICA, see e.g. [17]) on the gene expression experiment matrix to develop a low dimensional representation of the processes. They then build a minimal spanning tree (MST) on the distances developed from the resulting latent representation to reconstruct a Waddington’s landscape given by ICA. After some correction, if there are branching processes, they report the longest paths along the MST as the pseudo time backbone and the summed distances as the pseudo time ordering for each cell. This approach is called *Monocle.* However, this method relies on having rough estimates of the capture time to induce the ordering in the pseudo time estimate. Our probabilistic description of Waddington’s landscape relieves this requirement and allows for post analysis of data sets which do not provide such estimates.

Other methods apply deterministic non-linear dimensionality reductions and attempt to recover the underlying pseudo time in a probabilistic framework [6].

*Wishbone* [25] applies the t-SNE [32] algorithm to reduce the dimensionality of the expression matrix and then proceeds by averaging over *k*-nearest-neighbour graph ensembles to extract pseudo times.

If other methods correct for distances in the extracted landscape, they usually employ heuristics or additional data about capture times. For this, they rely on Euclidean distances between cells to overlay the extraction method of pseudo time (usually graphs, on which to go along). For us, we can employ non Euclidean distances in the landscape, following the topography of the probabilistic landscape to use in the graph. This stabilises the extraction of time along the graph. Outliers can create “short cuts” in the graph structure, which will be identified by the landscape’s topography.

A Riemannian geometry distorts distances, just as in a real map movement is not equally easy in all directions (it is easier to go down hill or follow roads) the underlying Waddington landscape has a topology which should be respected. Topslam landscapes are both non-linear and probabilistic and we correct, locally, for Riemannian distortions introduced by the non-linear surfaces. In the next section we will show how the combination of these three characteristics allows us to recover pseudo time *without* reliance on additional data or additional (correctional) algorithms for graph extraction, to correct for the underlying dimensionality reduction technique used.

In summary, we introduce a probabilistic approach to inferring Waddington landscapes and we consider the topological constraints of that landscape. In the next section we show how this idea can be used to improve pseudo time recovery for single cell data.

## 2 Application: Pseudo Time Recovery

Single cell gene expression experiments provide unprecedented access to the underlying processes and intrinsic functional relationships of and between cells. However, looking at single cells the extracted gene expression is prone to the heterogeneous variability from cell-to-cell. Such noise is not only technical (such as low amounts of RNA, dropout events etc. [19]), but also biological in origin (heterogeneity between cells of the same type).

Each cell is a functioning member of a local community of cells. Biology is based on an evolutionary progression, in which old systems are usually kept in place, when new ones are found. This introduces a lot of redundancies in such processes and makes extraction of information and evidence complex. Therefore, we use dimensionality reduction techniques to optimise and visualise the underlying landscape of the biological process.

Epigenetic progression is a discrete process that Waddington suggested could be visualised as part of a continuous landscape. However, the relationship between location on the landscape and the measured state of the cell is unlikely to be *linear.*

Further, when mapping natural landscapes, a laborious process of triangulation through high precision measurements is used to specify the map accurately. In the epigenetic landscape, no such precision is available. As a result it is vital that we sustain an estimate of our *uncertainty* about the nature of the landscape as we develop the map.

### 2.1 Simulation and Validation

Simulation was done by simulating 5 differentiation patterns of cells (Fig. 2). We then extracted pseudo time orderings of the cells in the simulation from 10 repetitions of creating gene expression measurements driven by the simulated differentiation patters (details Supplementary S1).

**Figure 2.**
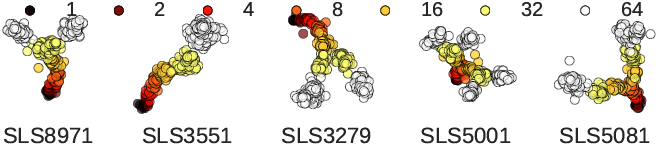
Simulated differentiation processes along cell stages. The cell stages are coloured from 1 to 64 cell stage and each simulation has its associated unique seed printed underneath. The selection of differentiation processes was done by visual inspection, strafing for variety and non overlapping profiles, so that a 2 dimensional landscape was possible.

#### 2.1.1 Simulation Results

We compare extracted pseudo time orderings for four methods in Table 1. The four methods we compare are Monocle [31], Wishbone [25], Bayesian GPLVM, and topslam. Shown are the linear regression correlation coefficients *ρ* and standard deviations over 30 tries between simulated and extracted time lines. From the simulation studies we can extract, that we can fully reconstruct the simulated time at an average correlation of approximately 90%[±7%] (Table 1). This is about 5% higher correlation then the next best method Wishbone (at 86%[±9%]). The construction of Waddington’s landscape ensures an improvement over the other methods in all simulated latent spaces, even if the intrinsic signal structure suits the other methods. Additionally, the consistency of our result is higher across the experiments, providing more reliable results over multiple experiments.

**Table 1.**
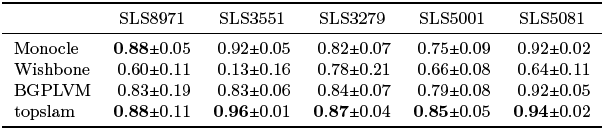
Simulation study results for gene expression matrices generated from simulated Waddington landscapes along a time line. Shown are linear regression Pearson correlation coefficients *ρ* between extracted and simulated time lines. Data sets where simulated from different differentiation profiles as described in Section S1.

The simulation results show that topslam is robust to repetition and differences in underlying surfaces, whereas the other methods fail in certain circumstances, especially when the underlying differentiation process gets complex (more branching). Thus, it is crucial to account for the topography of the dimensionality reduction technique, before extracting intrinsic signals (such as pseudo time) from the rich phenotypic characterisations of cells.

We also show, that we can use topslam to overlay a probabilistic Waddington’s landscape over the other dimensionality reduction techniques. This enables a corrected extraction of pseudo time estimates. This correction is shown to be never detrimental and can increase the correlation between extracted and simulated pseudo times (Supplementary S1). The supplementary material also contains results for a range of other dimensionality reduction techniques.

#### 2.1.2 Running Time

Our probabilistic treatment of landscape recovery and our principled correction of the topology mean that topslam is the slowest of the three approaches. The other two methods only apply heuristic corrections, gaining speed in the estimation of intrinsic signal ordering. Topslam averages at approximately 230*s* of run time to learn the landscape for the simulated 400 – 500 cells. (The number of genes does not play a significant role during optimisation, because of pre-computation of the empirical covariance matrix.) Wishbone averages at approximately 40*s* and Monocle at only 5*s*. However, as we’ve seen this faster compute comes at the expense of a significant loss of both accuracy and consistency. We now turn to deployment of topslam on real data.

## 3 Pseudo Time Extraction Mouse Cells

In this section we explore the performance of topslam on to real single cell qPCR (Supplementary S2.1). This shows the ability for topslam to extract intrinsic signals for existing and difficult single cell profiling techniques, which can bare difficulties because of high noise corruption and systematic errors (dropouts, detection limit etc.).

### 3.1 Mouse Embryonic Development Landscape

We extract the pseudo time for a mouse embryonic single cell qPCR experiment [13,19] of 437 cells, captured from one to 64 cell-state. In this experiment 48 genes where captured. We learn a landscape for the cells progression along time, capturing the differentiation process. The landscape then defines the progression of time by following the valleys of the topography, depicted in Figure 3.

**Figure 3.**
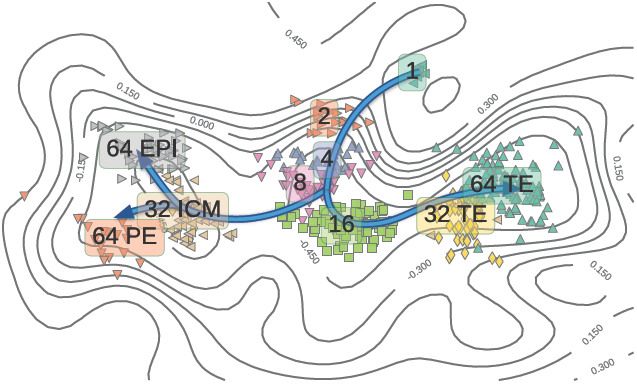
Representation of the probabilistic Waddington’s landscape. The contour lines represent heights of the landscape. The lower the landscape, the less “resistance” there is to move around. The time is then extracted along the cells such that it follows the landscape, depicted as splitting arrows. This also reflects the separate cell fates in the epigenetic progression of the cells.

Extracting the progression landscape from a qPCR single cell gene expression experiment [14] reveals the time line for the single cell progression in fine grained detail. We extract the landscape for the developmental cells and compute distances along the landscape through an embedded graph.

The starting cell needs to be given, whereas no more information is needed to extract the progression of (pseudo-) time along the graph. It is recommended to provide a leaf node in the graph, to ensure only one direction of time along Waddington’s landscape. If a cell in the middle is specified, the time will go positive in two directions starting from the starting cell. Thus, a leaf node will ensure, that the time runs in only one direction.

We can now use the extracted time to infer differing progression of gene expression through the progression of cells. In this particular data set we have a differentiation progress at hand, cells differentiating into three different cell states in the 64 cell stage: trophectoderm (TE), epiblast (EPI), and primitive endoderm (PE).

We use the same labelling of Guo *et al.* [14], which introduces some systematic errors (as explained in Section 1.1). With this differentiation, we can now plot gene expression along the timeline, revealing the dynamics of gene expression during differentiation and elucidating differentiation processes within different cell types (Fig. S9). Using the extracted pseudo time for different pathways in the cell stages, we can elucidate the differentiation process along time. We perform differential gene expression detection in time series experiments (e.g. [26]), and use the top ten differentially expressed genes as marker genes for the three cell stages (Table 2). We compiled the list as a comparison between stages, thus if a gene is duplicated in the comparison of stages it is a marker gene for the differentiation of the one stage from the two others (see e.g. for TE *Id2, Tspan8).* The differentiation takes place in the 16 and 32 cell stages (Figure 4). Having the time series as differential expressed marker genes, we can plot the exact time line of when genes get differentially expressed along pseudo time (Figure 4).

**Table 2.**
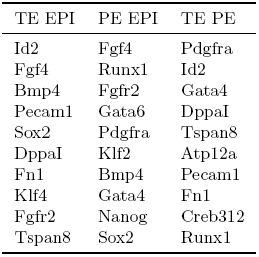
Marker genes for differentiation between the three cell stages compiled from time series differential expression along the pseudotime. Shown are the ten most differentially expressed genes, pairwise between the three stages. For example *Id2* is differentially expressed between (TE and EPI) and between (TE and PE). This means it is a marker gene for TE, as it behaves differently from the two other differentiation stages, but not within the two others. *Id2* is known to be a marker for TE.

| TE EPI | PE EPI | TE PE |
| --- | --- | --- |
| Id2 | Fgf4 | Pdgfra |
| Fgf4 | Runx1 | Id2 |
| Bmp4 | Fgfr2 | Gata4 |
| Pecam1 | Gata6 | Dppa1 |
| Sox2 | Pdgfra | Tspan8 |
| Dppa1 | Klf2 | Atp12a |
| Fn1 | Bmp4 | Pecam1 |
| Klf4 | Gata4 | Fn1 |
| Fgfr2 | Nanog | Creb312 |
| Tspan8 | Sox2 | Runx1 |

**Figure 4.**
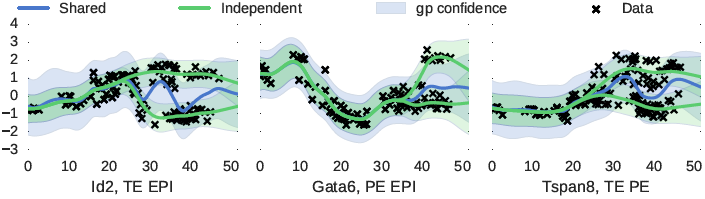
Some example plots for the marker gene extraction. In green you can see the individual fits of two GPs, sharing one prior, and in blue the shared fit of one GP to all the data. Differential expression is decided on which of those two models (green or blue) fits the data better. Note the time line elucidates when (in time) the gene can be used as a marker gene. *Gata6* is a known marker for TE, but evidently it is also differentially expressed in mice between PE and EPI differentiation states.

Comparison with results using other dimensionality reduction techniques, show that the other methods are not able to capture the non-linearities in the data (topslam is our method, Figure 5). We can also see the representation of Waddington’s landscape as shaded area, we want to stay in light areas.

**Figure 5.**
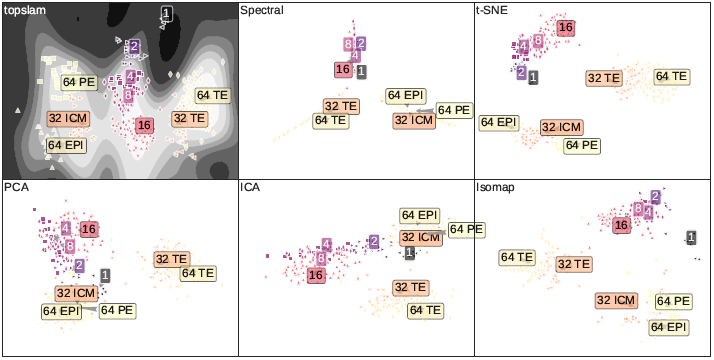
Comparison plots between different dimensionality reduction techniques for the Guo *et al.* data set of developmental mouse embryonic stem cells [14]. As can be seen, only topslam (probabilistic Waddington landscape) can fully identify the relationships between the cells and order them correctly for pseudo time extraction. t-SNE is the underlying method Wishbone relies on and ICA the one for Monocle.

Using the probabilistic interpretation of Waddington’s landscape as a correction for the embedding and extraction techniques, we can extract pseudo time information more clearly and without additional information to ensure the time line extracted corresponds to the cell stages as seen in Guo *et al.* [14].

## 4 Conclusion

We have introduced a probabilistic approach to inferring Waddington landscapes. We use rich phenotype information to characterise the landscape and probabilistic inference techniques to infer a non-linear mapping from the landscape to the phenotype. Our approach allows us to respect the topology of the landscape when extracting distances and we show the advantages of this idea when reconstructing pseudo times from single cell data. Summarising single cells in this manner represents a powerful approach for understanding the evolution of their genetic profile, a critical stage in understanding development and cancer.

## 5 Methods

### 5.1 Data

For description of single cell transcriptome extraction techniques please refer to supplementary material S2.

### 5.2 Code

A package topslam written in python (based on GPy [12]) is provided for users to apply the methods described in this work.

https://github.com/mzwiessele/topslam

We supply all topslam correction methods in this package, including different graph extraction techniques. Additionally, we supply optimisation routines for the dimensionality reduction technique. For you convenience we include plotting routines and data filtering methods alongside the package.

### 5.3 Extracting Pseudo Time

The most common way of extracting pseudo time orderings is done with the following stages:

1. Extract lower dimensional representation 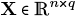 of gene expression matrix 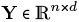 with *n* samples as rows and *d* genes as columns. The lower dimensional representation is often chosen to have *q* = 2 dimensions, as the dimensionality reduction techniques do not express a selection criterion and two dimensions are convenient for visualisation.
2. Supply starting point **s** ∈ **X** of pseudo time ordering extracted in the next step.
3. Extract distance information about cells by following the landscape by a graph structure, sometimes a tree, or k-nearest-neighbour graph.
4. Extract the ordering of cells along the graph structure extracted in the above step (including smoothing, branch detection, and/or clustering).

#### 5.3.1 Topslam Approach

Standard approaches each miss at least one important component of the mapping problem. Monocle assumes a linear map, a highly unrealistic assumption.

Wishbone [25] makes use of a non-linear method but does not consider the *topography* of the map when developing pseudo time orderings. The topography of the epigenetic landscape influences distances between cells on the landscape, and therefore their effective relative positions to each other.

Our approach, a topologically corrected simultaneous localisation and mapping of cells, topslam, proposes to make use of a *probabilistic non-linear* dimensionality reduction technique, also used in many other single cell transcriptomics applications [3-5,22]. The probabilistic nature of the dimensionality reduction technique is used for extracting the Waddington landscapes *with* associated uncertainties. Further, we are able to take account of the local topography when extracting pseudo times, correcting distances by applying non Euclidean metrics along the landscape [30].

To perform pseudo time extraction with topslam we build a minimum spanning tree (or k-nearest-neighbour graph) along the latent landscape uncovered by topslam. This allows the spanning tree to naturally follow the landscape topography and makes any corrections post extraction obsolete. For a more detailed description of the approach see supplementary material [S3,S4].

## Acknowledgements

The authors would like to thank Aleksandra Kołodziejczyk, Alexis Boukouvalas, Florian Büttner, Sarah Teichmann, and Magnus Rattray for useful discussion and clarifying correspondence.

## References

1. S. C. Bendall, K. L. Davis, E.-a. D. Amir, M. D. Tadmor, E. F. Simonds, T. J. Chen, D. K. Shenfeld, G. P. Nolan, and D. Peer. Single-cell trajectory detection uncovers progression and regulatory coordination in human B cell development. Cell, 157(3):714–725, 2014.

2. S. Bhattacharya, Q. Zhang, and M. E. Andersen. A deterministic map of Waddington’s epigenetic landscape for cell fate specification. BMC systems biology, 5(1):85, 2011.

3. F. Buettner, V. Moignard, B. Göttgens, and F. J. Theis. Probabilistic PCA of censored data: accounting for uncertainties in the visualization of high-throughput single-cell qPCR data. Bioinformatics, 30(13):1867–1875, 2014.

4. F. Buettner, K. N. Natarajan, F. P. Casale, V. Proserpio, A. Scialdone, F. J. Theis, S. A. Teichmann, J. C. Marioni, and O. Stegle. Computational analysis of cell-to-cell heterogeneity in single-cell RNA-sequencing data reveals hidden subpopulations of cells. Nature biotechnology, 33(2):155–160, 2015.

5. F. Buettner and F. J. Theis. A novel approach for resolving differences in single-cell gene expression patterns from zygote to blastocyst. Bioinformatics, 28(18):i626–i632, 2012.

6. K. Campbell and C. Yau. Bayesian Gaussian Process Latent Variable Models for pseudotime inference in single-cell RNA-seq data. bioRxiv, page 026872, 2015.

7. P. Dalerba, T. Kalisky, D. Sahoo, P. S. Rajendran, M. E. Rothenberg, A. A. Leyrat, S. Sim, J. Okamoto, D. M. Johnston, D. Qian, et al. Single-cell dissection of transcriptional heterogeneity in human colon tumors. Nature biotechnology, 29(12):1120–1127, 2011.

8. A. Diaz, S. J. Liu, C. Sandoval, A. Pollen, T. J. Nowakowski, D. A. Lim, and A. Kriegstein. Scell: integrated analysis of single-cell rna-seq data. Bioinformatics, page btw201, 2016.

9. B. Ferris, D. Fox, and N. D. Lawrence. WiFi-SLAM Using Gaussian Process Latent Variable Models. In IJCAI, volume 7, pages 2480–2485, 2007.

10. R. A. Gibbs, J. W. Belmont, P. Hardenbol, T. D. Willis, F. Yu, H. Yang, L.-Y. Ch’ang, W. Huang, B. Liu, Y. Shen, et al. The international HapMap project. Nature, 426(6968):789–796, 2003.

11. A. Git, H. Dvinge, M. Salmon-Divon, M. Osborne, C. Kutter, J. Hadfield, P. Bertone, and C. Caldas. Systematic comparison of microarray profiling, real-time PCR, and next-generation sequencing technologies for measuring differential microRNA expression. RNA, 16(5):991–1006, May 2010.

12. GPy. GPy: A Gaussian process framework in python. http://github.com/SheffieldML/GPy, since 2012.

13. D. Grün and A. van Oudenaarden. Design and analysis of single-cell sequencing experiments. Cell, 163(4):799–810, 2015.

14. G. Guo, M. Huss, G. Q. Tong, C. Wang, L. Li Sun, N. D. Clarke, and P. Robson. Resolution of cell fate decisions revealed by single-cell gene expression analysis from zygote to blastocyst. Developmental cell, 18(4):675–685, 2010.

15. D. S. Horner, G. Pavesi, T. Castrignanò, P. D. De Meo, S. Liuni, M. Sammeth, E. Picardi, and G. Pesole. Bioinformatics approaches for genomics and post genomics applications of next-generation sequencing. Brief Bioinform, 11(2):181–97, Mar 2010.

16. S. Huang. The molecular and mathematical basis of waddington’s epigenetic landscape: A framework for post-darwinian biology? Bioessays, 34(2):149–157, 2012.

17. A. Hyvärinen, J. Karhunen, and E. Oja. Independent component analysis, volume 46. John Wiley & Sons, 2004.

18. S. Islam, U. Kjällquist, A. Moliner, P. Zajac, J.-B. Fan, P. Lönnerberg, and S. Linnarsson. Characterization of the single-cell transcriptional landscape by highly multiplex RNA-seq. Genome research, 21(7):1160–1167, 2011.

19. T. Kalisky and S. R. Quake. Single-cell genomics. Nat Meth, 8(4):311–314, Apr. 2011.

20. C. Marr, J. X. Zhou, and S. Huang. Single-cell gene expression profiling and cell state dynamics: collecting data, correlating data points and connecting the dots. Current opinion in biotechnology, 39:207–214, 2016.

21. A. McDavid, G. Finak, P. K. Chattopadyay, M. Dominguez, L. Lamoreaux, S. S. Ma, M. Roederer, and R. Gottardo. Data exploration, quality control and testing in single-cell qPCR-based gene expression experiments. Bioinformatics, 29(4):461–467, 2013.

22. V. Moignard, I. C. Macaulay, G. Swiers, F. Buettner, J. Schütte, F. J. Calero-Nieto, S. Kinston, A. Joshi, R. Hannah, F. J. Theis, et al. Characterization of transcriptional networks in blood stem and progenitor cells using high-throughput single-cell gene expression analysis. Nature cell biology, 15(4):363–372, 2013.

23. P. Paschou, E. Ziv, E. G. Burchard, S. Choudhry, W. Rodriguez-Cintron, M. W. Mahoney, and P. Drineas. PCA-correlated SNPs for structure identification in worldwide human populations. PLoS Genet, 3(9):1672–86, Sep 2007.

24. C. Rampon, C. H. Jiang, H. Dong, Y.-P. Tang, D. J. Lockhart, P. G. Schultz, J. Z. Tsien, and Y. Hu. Effects of environmental enrichment on gene expression in the brain. PNAS, November 2000.

25. M. Setty, M. D. Tadmor, S. Reich-Zeliger, O. Angel, T. M. Salame, P. Kathail, K. Choi, S. Bendall, N. Friedman, and D. Pe’er. Wishbone identifies bifurcating developmental trajectories from single-cell data. Nature Biotechnology, 2016.

26. O. Stegle, K. J. Denby, E. J. Cooke, D. L. Wild, Z. Ghahramani, and K. M. Borgwardt. A robust Bayesian two-sample test for detecting intervals of differential gene expression in microarray time series. Journal of Computational Biology, 17(3):355–367, 2010.

27. S. Thrun and J. J. Leonard. Simultaneous localization and mapping. In Springer handbook of robotics, pages 871–889. Springer, 2008.

28. M. E. Tipping and C. M. Bishop. Mixtures of probabilistic principal component analyzers. Neural computation, 11(2):443–482, 1999.

29. M. K. Titsias and N. D. Lawrence. Bayesian Gaussian Process Latent Variable Model. Artificial Intelligence and Statistics, 2010.

30. A. Tosi, S. Hauberg, A. Vellido, and N. D. Lawrence. Metrics for probabilistic geometries. In Proceedings of 30th Conference on Uncertainty in Artificial Intelligence (uai 2014). AUAI Press Corvallis, 2014.

31. C. Trapnell, D. Cacchiarelli, J. Grimsby, P. Pokharel, S. Li, M. Morse, N. J. Lennon, K. J. Livak, T. S. Mikkelsen, and J. L. Rinn. The dynamics and regulators of cell fate decisions are revealed by pseudotemporal ordering of single cells. Nature biotechnology, 32(4):381–386, 2014.

32. L. Van der Maaten and G. Hinton. Visualizing data using t-SNE. Journal of Machine Learning Research, 9(2579-2605):85, 2008.

33. C. Waddington. Principles of development and differentiation, CH Waddington. Current concepts in biology series., 1966.

34. C. H. Waddington. The strategy of the genes, volume 20. Routledge, 2014.

35. W. Wu and J. Wang. Potential and flux field landscape theory. i. global stability and dynamics of spatially dependent non-equilibrium systems. The Journal of chemical physics, 139(12):121920, 2013.

36. J. X. Zhou, M. Aliyu, E. Aurell, and S. Huang. Quasi-potential landscape in complex multi-stable systems. Journal of the Royal Society Interface, 9(77):3539–3553, 2012.

